# PISCES: a package for rapid quantitation and quality control of large scale mRNA-seq datasets

**DOI:** 10.1101/2020.12.01.390575

**Authors:** Matthew D. Shirley, Viveksagar K. Radhakrishna, Javad Golji, Joshua M. Korn

## Abstract

PISCES eases processing of large mRNA-seq experiments by encouraging capture of metadata using simple textual file formats, processing samples on either a single machine or in parallel on a high performance computing cluster (HPC), validating sample identity using genetic fingerprinting, and summarizing all outputs in analysis-ready data matrices. PISCES consists of two modules: 1) compute cluster-aware analysis of individual mRNA-seq libraries including species detection, SNP genotyping, library geometry detection, and quantitation using salmon, and 2) gene-level transcript aggregation, transcriptional and read-based QC, TMM normalization and differential expression analysis of multiple libraries to produce data ready for visualization and further analysis.

PISCES is implemented as a python3 package and is bundled with all necessary dependencies to enable reproducible analysis and easy deployment. JSON configuration files are used to build and identify transcriptome indices, and CSV files are used to supply sample metadata and to define comparison groups for differential expression analysis using DEseq2. PISCES builds on many existing open-source tools, and releases of PISCES are available on GitHub or the python package index (PyPI).

## Introduction

Since the first description of RNA-seq [1], methods for RNAseq quantification have rapidly increased in sensitivity and decreased in required processing time. Recent improvements in speed have been achieved by removing the step of full read alignment to a reference [2][3][4]. While these “alignment-free” (also known as “pseudoalignment,” “quasi-mapping,” or “lightweight alignment”) methods have achieved real speed gains, this comes at a loss of compatibility with existing RNAseq workflows, including important integrity checks. Alignment-free quality control (QC) and sample identification methods for RNAseq libraries are not readily available, and existing packages [5] for QC are incompatible with the data formats produced by these new quantification methods. Decreased sequencing cost have also led to higher throughput of sequencing, with an associated desire for faster quantification and turnaround of primary analysis. Typical RNAseq experiments may now be composed of hundreds to thousands of individual samples, each with descriptive variables for downstream analysis, which increases the complexity of data and metadata management. Increased number of samples presents more opportunities for sample mixups in the lab. Complex designs or cross-experiment meta-analyses further present opportunities for in silico label swapping. Tracking of these crucial metadata are usually left to the sequencing facility or individual analyst. With no widely agreed upon standards for metadata exchange, information about RNAseq libraries may be lost.

With these types of experiments in mind, we built PISCES with an alignment-free human SNP fingerprinting method for checking sample identity, efficient FASTQ QC, and novel QC of the transcriptomic and genomic compartments without the need for full genomic alignment. PISCES is driven by a well-defined CSV metadata format (Table 1), which limits opportunities for label swaps and encourages analysts to retain a minimal set of metadata necessary to reproduce an analysis.

**Table 1:** Experimental metadata such as treatment variables and covariates of interest (cell line, time point etc.) are described in CSV format.

| Directory | SRA | SampleID | Treatment | Genotype | CellLine |
| --- | --- | --- | --- | --- | --- |
| SRR5024105 | SRR5024105 | GSM2392606 | Vehicle | Wildtype | MCF7 |
| SRR5024106 | SRR5024106 | GSM2392607 | Vehicle | Wildtype | MCF7 |
| SRR5024107 | SRR5024107 | GSM2392608 | Vehicle | Wildtype | MCF7 |
| SRR5024108 | SRR5024108 | GSM2392609 | Vehicle | Wildtype | MCF7 |
| SRR5024109 | SRR5024109 | GSM2392610 | Vehicle | Y537S | MCF7 |
| SRR5024110 | SRR5024110 | GSM2392611 | Vehicle | Y537S | MCF7 |
| SRR5024111 | SRR5024111 | GSM2392612 | Vehicle | Y537S | MCF7 |
| SRR5024112 | SRR5024112 | GSM2392613 | Vehicle | Y537S | MCF7 |
| SRR5024113 | SRR5024113 | GSM2392614 | Vehicle | D538G | MCF7 |
| SRR5024114 | SRR5024114 | GSM2392615 | Vehicle | D538G | MCF7 |
| SRR5024115 | SRR5024115 | GSM2392616 | Vehicle | D538G | MCF7 |
| SRR5024116 | SRR5024116 | GSM2392617 | Vehicle | D538G | MCF7 |
| SRR5024117 | SRR5024117 | GSM2392618 | Estrogen | Wildtype | MCF7 |
| SRR5024118 | SRR5024118 | GSM2392619 | Estrogen | Wildtype | MCF7 |
| SRR5024119 | SRR5024119 | GSM2392620 | Estrogen | Wildtype | MCF7 |
| SRR5024120 | SRR5024120 | GSM2392621 | Estrogen | Wildtype | MCF7 |
| SRR5024121 | SRR5024121 | GSM2392622 | Estrogen | Y537S | MCF7 |
| SRR5024122 | SRR5024122 | GSM2392623 | Estrogen | Y537S | MCF7 |
| SRR5024123 | SRR5024123 | GSM2392624 | Estrogen | Y537S | MCF7 |
| SRR5024124 | SRR5024124 | GSM2392625 | Estrogen | Y537S | MCF7 |
| SRR5024125 | SRR5024125 | GSM2392626 | Estrogen | D538G | MCF7 |
| SRR5024126 | SRR5024126 | GSM2392627 | Estrogen | D538G | MCF7 |
| SRR5024127 | SRR5024127 | GSM2392628 | Estrogen | D538G | MCF7 |
| SRR5024128 | SRR5024128 | GSM2392629 | Estrogen | D538G | MCF7 |

In addition to streamlining primary analysis (transcript and gene quantification) at the single sample level, PISCES provides methods for executing experiment-level analyses. One common secondary analysis method for RNAseq libraries is differential gene expression analysis (DGE). PISCES implements DGE using a wrapper for DEseq2 [6], with contrast groups defined using descriptive variables referenced in the CSV metadata file. PISCES creates summarized data matrices of transcript-level counts and TPM, as well as gene-level counts, TPM, and log_2_ fold-change calculated using the median of a user-specified reference group for each condition. Trimmed mean of M-values normalization [7] as implemented in edgeR is used to adjust TPM abundances of protein coding genes to account for differences in RNAseq library transcript composition, e.g. a varying amount of non-coding transcripts due to pre-mRNA contamination or incomplete rRNA depletion.

Finally, with increasing numbers of RNAseq libraries comes an increased computational burden. PISCES communicates directly with modern compute clusters using the distributed resource management application API (DRMAA) to efficiently submit and monitor jobs processing individual RNAseq libraries.

## Implementation of the PISCES package

PISCES is provided as a Python 3 package, including all tools and dependencies, which users can easily install using common packaging tools such as pip on modern Linux and MacOS machines.

Versioned releases are available from both Github and PyPI. Portions of PISCES are implemented in the R statistical computing language, and these dependencies, as well as other binary dependencies, are also bundled and installed automatically using renv [8]. Workflows can be run either locally, on a single machine, or on a compute cluster supporting the DRMAA interface. Individual RNAseq libraries may be analyzed without associated metadata by passing FASTQ files directly into PISCES, or entire experiments may be defined using a CSV format metadata file containing, at a minimum: sample names (*SampleID*), FASTQ file locations (*Fastq1 and optionally Fastq2*) or NCBI SRA run accessions (*SRA*), and output directories for each sample (*Directory*). The metadata file may also contain any descriptive information associated with the biological samples such as treatment, batch, or physical sample QC metrics. This descriptive information can be used by PISCES to identify sample groupings and reference samples for calculating normalized log2 fold changes, and to identify groups of samples for generating QC and exploratory figures. Differential expression is performed using DEseq2, using contrasts defined in a separate CSV format (Table 2) that describes covariates of interest referenced in the metadata CSV file. Transcriptomic QC metrics are calculated from the ratios of unprocessed intronic transcripts, mature processed transcripts, and intergenic regions. Several library QC metrics are inferred using only kmer counting directly from FASTQ files, including species detection, strand detection, and human SNP fingerprint for sample identification.

**Table 2:**
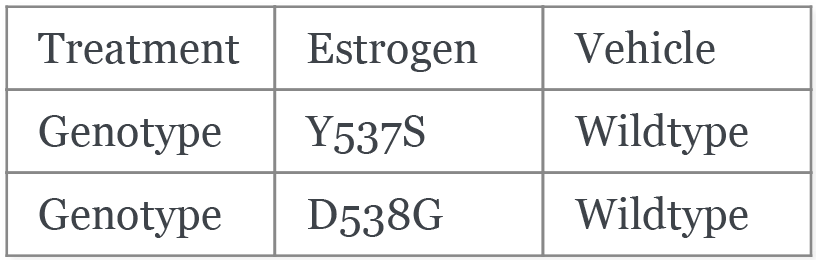
Contrasts for differential expression analysis are defined using the variables present in the metadata table. Columns in this file do not have headers, but correspond to `covariate name, experimental value, control value`.

### The PISCES workflow

The PISCES workflow (Fig. 1) is designed to be **flexible**, requiring minimal setup to describe an experiment; **scalable**, running on one to many samples either locally or on a compute cluster; **fast**, using alignment-free methods to generate QC metrics; and **reproducible**, driven by easily generated configuration files and versioned transcriptome annotation, with automated generation of transcriptome index files. We demonstrate each step of the PISCES workflow on a real experiment (Fig. 2) and show examples of PISCES outputs on real data.

**Fig. 1.**
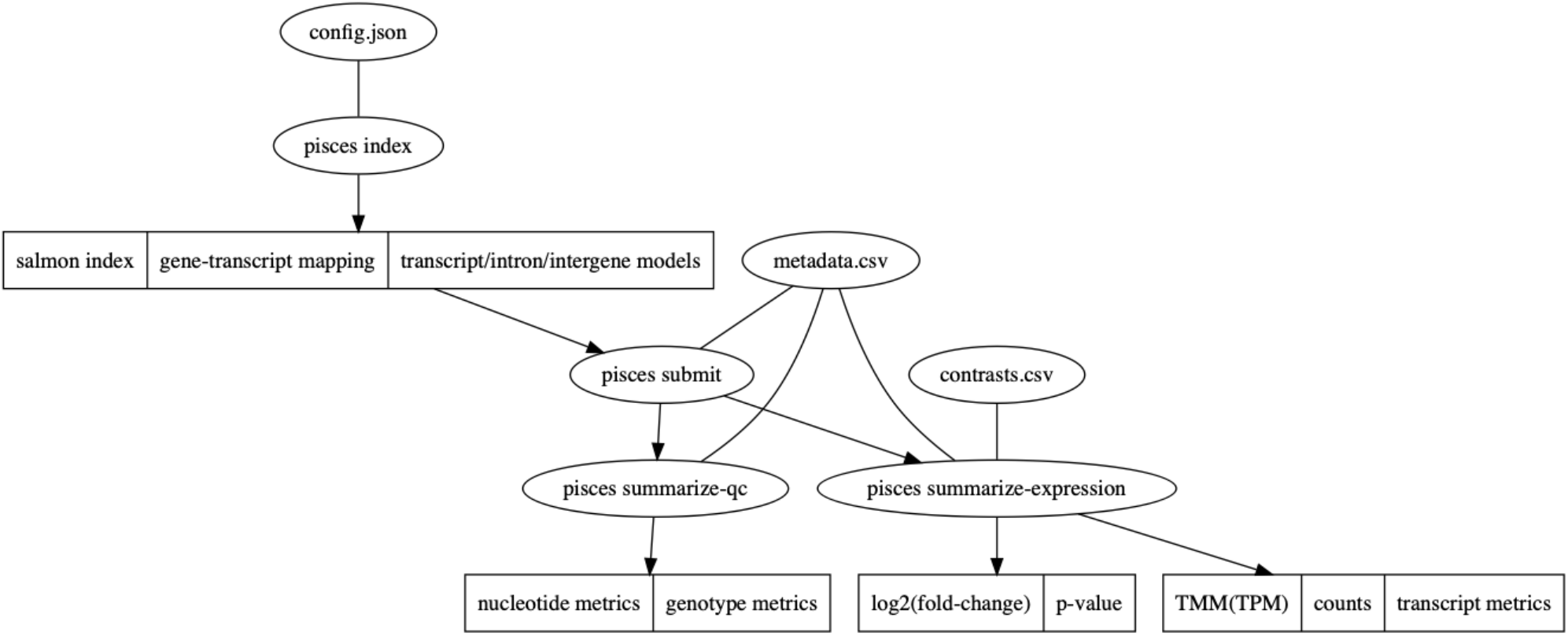
Overview of PISCES workflow, demonstrating configuration file inputs and descriptions of processes and outputs for each PISCES subcommand (index, submit, summarize-expression and summarize-qc).

**Fig. 2.**
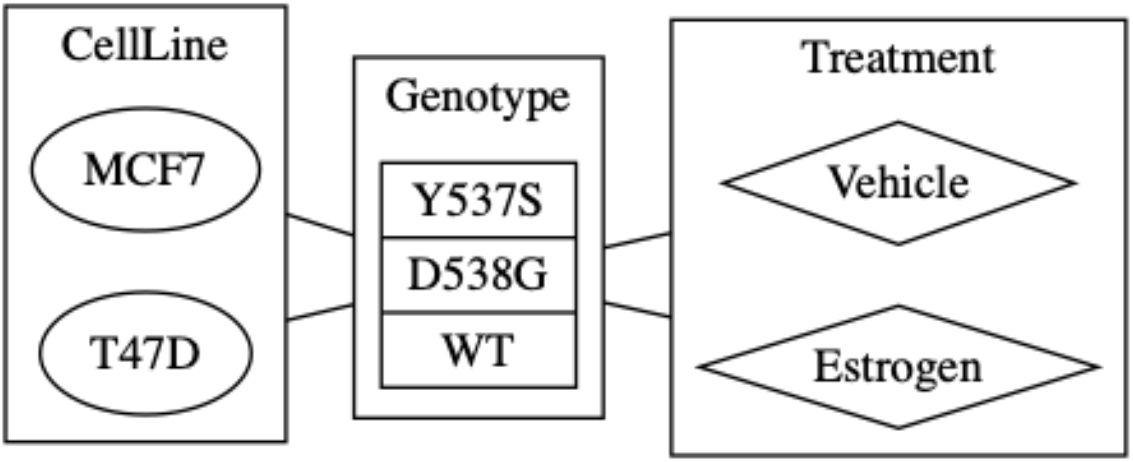
Treatment information for SRP093386

## Results

As a demonstration of the speed and simplicity of the PISCES workflow we reprocess an NCBI SRA study consisting of two breast cancer cell lines, MCF7 and T47D, with Y537S or D538G mutations introduced in the ESR1 (estrogen receptor) gene and treated with Vehicle or Estrogen (Fig. 2).

### pisces index builds custom transcriptome indices

PISCES is distributed with configuration for human, mouse, and human/mouse chimera transcriptomes (for xenograft experiments) derived from Gencode v32 and Gencode vM23 [9]. Custom transcriptomes can easily be defined in a JSON configuration file format. Extra sequences (such as bacterial genomes, spike ins for library prep, or genetic knock-ins in the experimental design) can be added to transcriptomes as arbitrary FASTA formatted files. PISCES builds transcript sequences using the GTF file format by first creating all defined transcript models per gene, and then concatenating all unique intron sequences to form one intronic transcript model per gene. In this way PISCES enables simultaneous quantification of both fully processed mRNA and contaminating pre-mRNA transcripts. Intergenic regions can also be captured, allowing quantification of all transcriptomic compartments of an organism’s genomic assembly. PISCES leverages the pufferfish index [10] enhancements in salmon version 1.0 and above which allows indexing of vastly larger transcript sequences.

### CSV formatted metadata files are input for PISCES

We include a metadata file describing libraries generated from MCF7 cell lines in [11] as a test script, distributed with PISCES. The following analysis, reproduced from this test script, serves as an example of a typical RNAseq workflow from FASTQ files to differential expression calls. In this example, PISCES uses the NCBI SRA toolkit to directly obtain sequencing data from NCBI servers, although paths to local data can be substituted in *Fastq1*, and optionally *Fastq2* columns.

With the above two files, an example PISCES workflow is only a few commands, and can be run easily on most Linux and MacOS machines.

~~~
$ pip install novartis-pisces
$ pisces index
$ pisces submit -m metadata.csv
$ pisces summarize-qc -m metadata.csv -f fingerprint.txt
$ pisces summarize-expression -m metadata.csv -f contrasts.csv -d “~Treatment + Genotype”
~~~

A formula for linear modeling of the experiment (**-d**) specifies the **Treatment** and **Genotype** covariates, and the interaction between the two, and p-values and log2 fold change are computed from the fit for each comparison defined in the contrasts CSV. The resulting summarized expression and QC data is output to tab separated text files suitable for further analysis and visualization using a suitable analytics environment such as Jupyter or RStudio.

### pisces submit runs PISCES on a DRMAA enabled HPC

The PISCES `pisces submit` command simplifies the task of executing the main PISCES workflow on a typical compute cluster, meaning a grid compute cluster using SGE, UGE, SLURM, Torque, or another DRMAA enabled job scheduler. The `pisces submit` command takes an appropriately formatted metadata file similar to Table 1 as input and dispatches multiple `pisces run` jobs, passing along any extra arguments a user supplies, as well as any cluster resource limits such as maximum job runtime, maximum memory, and number of CPU cores requested. *pisces submit* then monitors job log output and reports job status (number of jobs queued, running, finished), enabling a user to quickly process and track a large number of RNAseq libraries in parallel.

### pisces summarize-qc aggregates genetic QC metrics

Sample quality and identity is paramount for interpretable RNAseq results, but is missing from most alignment-free workflows. Here, read sequence quality metrics are assessed using fastqp version 0.3.4. For studies involving human samples, PISCES uses exact kmer counting (k=21) to generate a VCF per sample based on 226 chosen high minor-allele frequency SNPs. Log-odds scores representing the likelihood that two samples are genetically identical are used to generate a sample identity matrix. To demonstrate PISCES’ ability to match genetic identity even across independent experiments, we included breast cancer cell lines from the Cancer Cell Line Encyclopedia [12] with our example SRA analysis. In (Fig. 3) we show the genetic fingerprints of the CCLE [12] T47D and MCF7 cell lines match the expected clusters from the SRA experiment. Additionally, one can detect cell line pairs that were derived from the same patient (the KPL1 cell line which is a derivative of MCF7, and the autologous pair AU565 and SKBR3.)

**Fig. 3.**
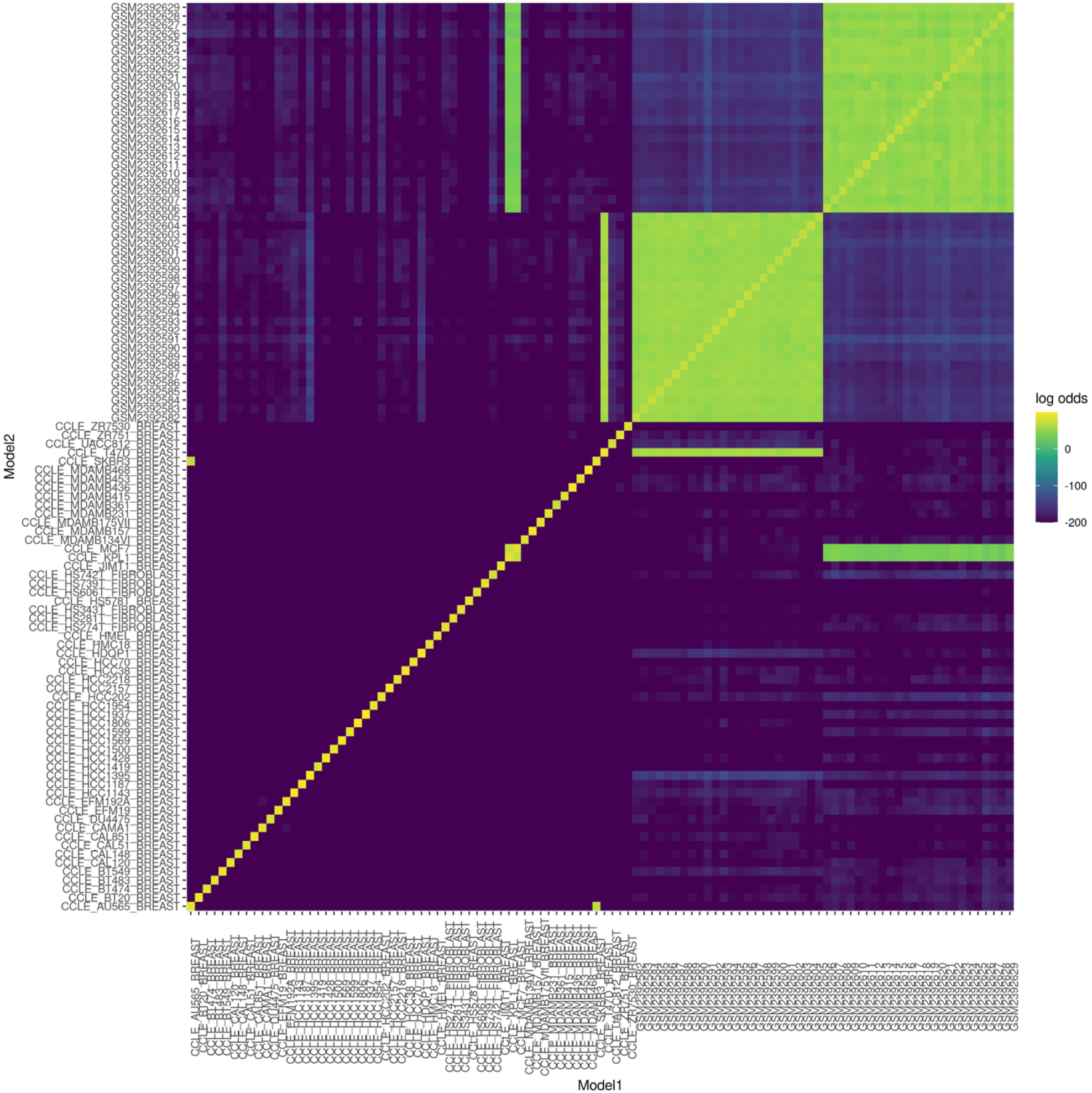
Sample identity matrix showing two clusters of samples, corresponding to libraries derived from MCF7 and T47D cell line models. The p_same score is a log-odds value where 0 indicates an exact genetic match and more negative values indicate a lower probability of genetic identity. Breast cancer cell lines from the publicly available CCLE are shown, demonstrating PISCES ability to integrate datasets across experiments.

### pisces summarize-expression aggregates transcript and gene abundance estimates

PISCES uses salmon [2] for estimating transcript and gene abundances from RNAseq libraries. Salmon is computationally efficient as well as accurate in assignment of reads to transcripts [13]. PISCES includes Salmon version 1.3.0. Library type parameters (strand-specific, read pairs, and read pair orientation) are inferred from input FASTQ files. Salmon is run with default parameters as well as specifying `--seqBias, --gcBias`, and `--useVBOpt`. The `pisces summarize-expression` subcommand aggregates transcriptomic counts and TPM estimates for multiple samples and summarizes these values to gene-level data matrices with appropriate column names defined in the metadata CSV file. TMM normalization is applied to the TPM estimates to control for highly expressed outlier transcripts. If unprocessed transcript models are specified during `pisces index`, data matrices for “intronic” and “intergene” transcriptomic compartments are output and a summary of percent coding, intergenic and intronic sequence estimates can be used to determine relative sample QC thresholds Fig. 4.

**Fig. 4.**
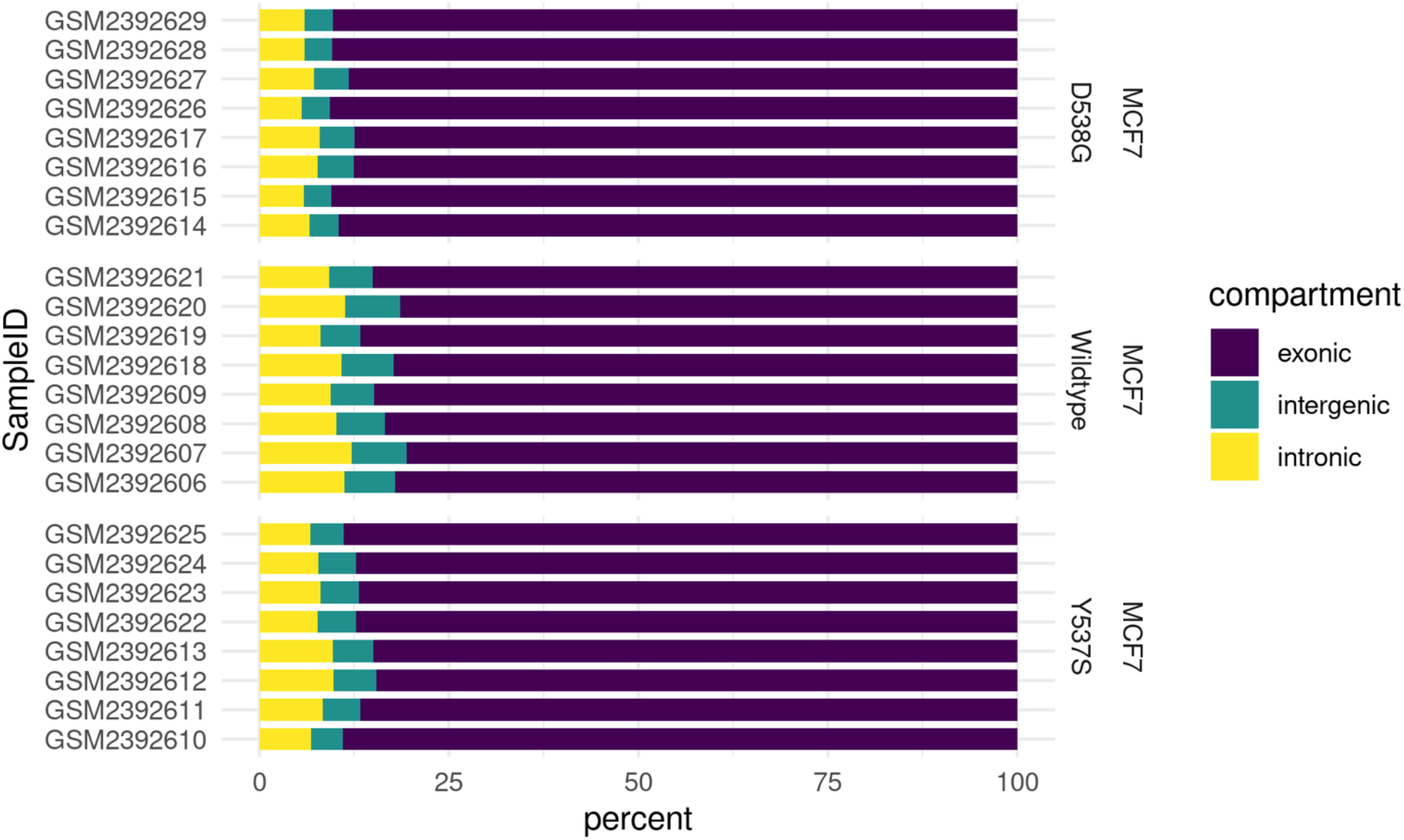
Percent coding, intronic, and intergenic read content is calculated salmon counts of processed transcript models, transcript models of introns, and intergenic sequences.

If contrasts (Table 2) and a formula are specified, differential gene expression is calculated and output to a tidy data table. These in turn can be visualized using standard graphing techniques, for example as a volcano plot Fig. 5.

**Fig. 5.**
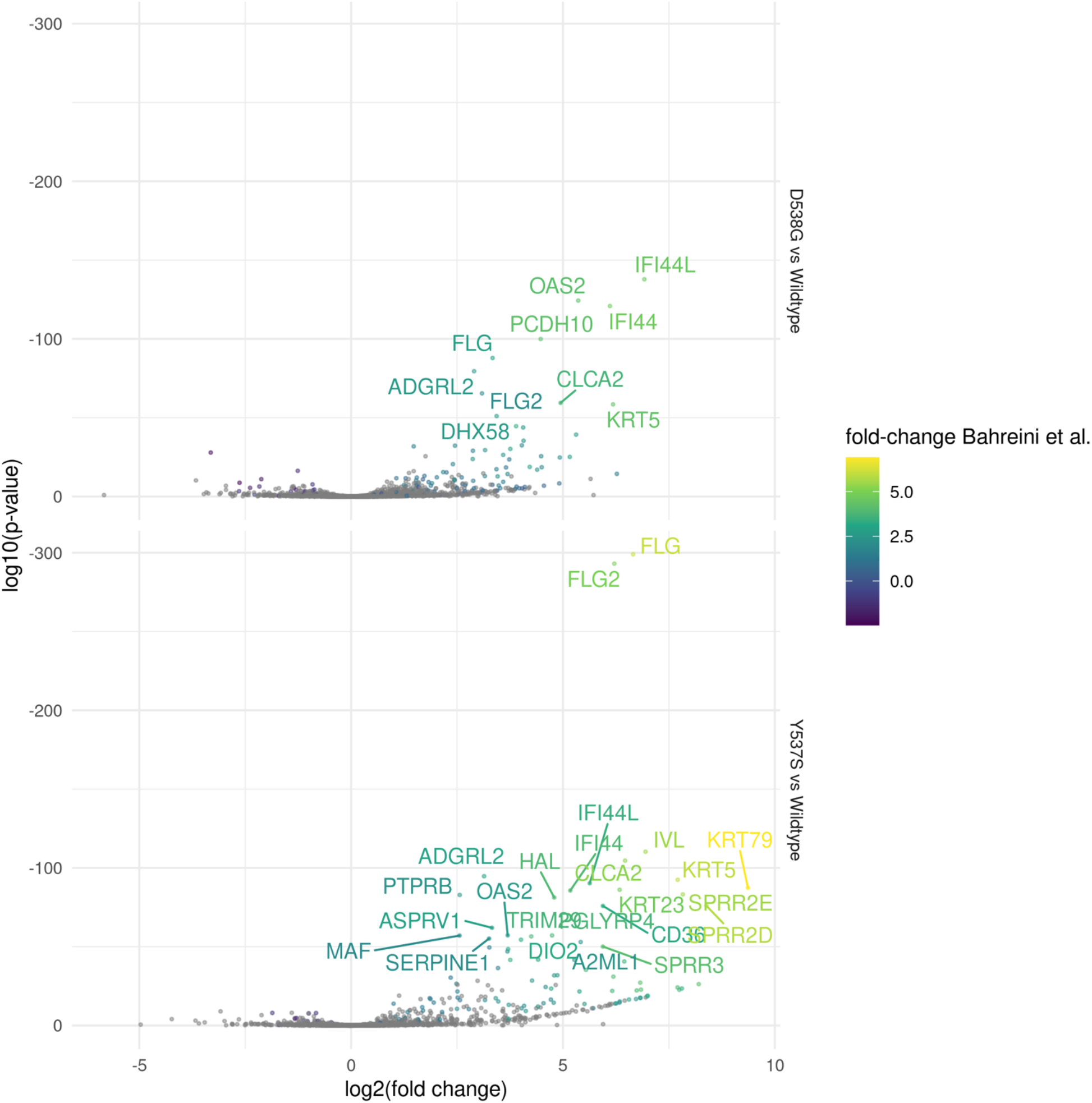
Volcano plot of differentially expressed genes (DEGs) identified by DESeq2, with **p < 0.01**. DEGs previously identified from [11] shown with the fold change stated in the original publication.

### pisces genomic indexing and repeat masking

During development of PISCES it became clear that the expression of certain genes was detected under biologically improbable contexts (such as aberrant immune cell marker gene expression in non-immune cancer cell culture). Upon closer inspection it appeared that many of these genes with unexpectedly high expression were mapping reads to repetitive regions of the transcript sequence. These repetitive regions are masked by RepeatMasker in the UCSC repeat masked genome assemblies, and so by default the pisces index command will hard-mask any soft-masked characters in the input genome assembly. The effect of repeat masking on salmon TPM estimates is that many genes with apparently low-to-medium abundance are significantly reduced, indicating their TPM estimates were driven primarily by repeat sequences and were likely a result of incorrectly assigned reads (Fig. 6).

**Fig. 6.**
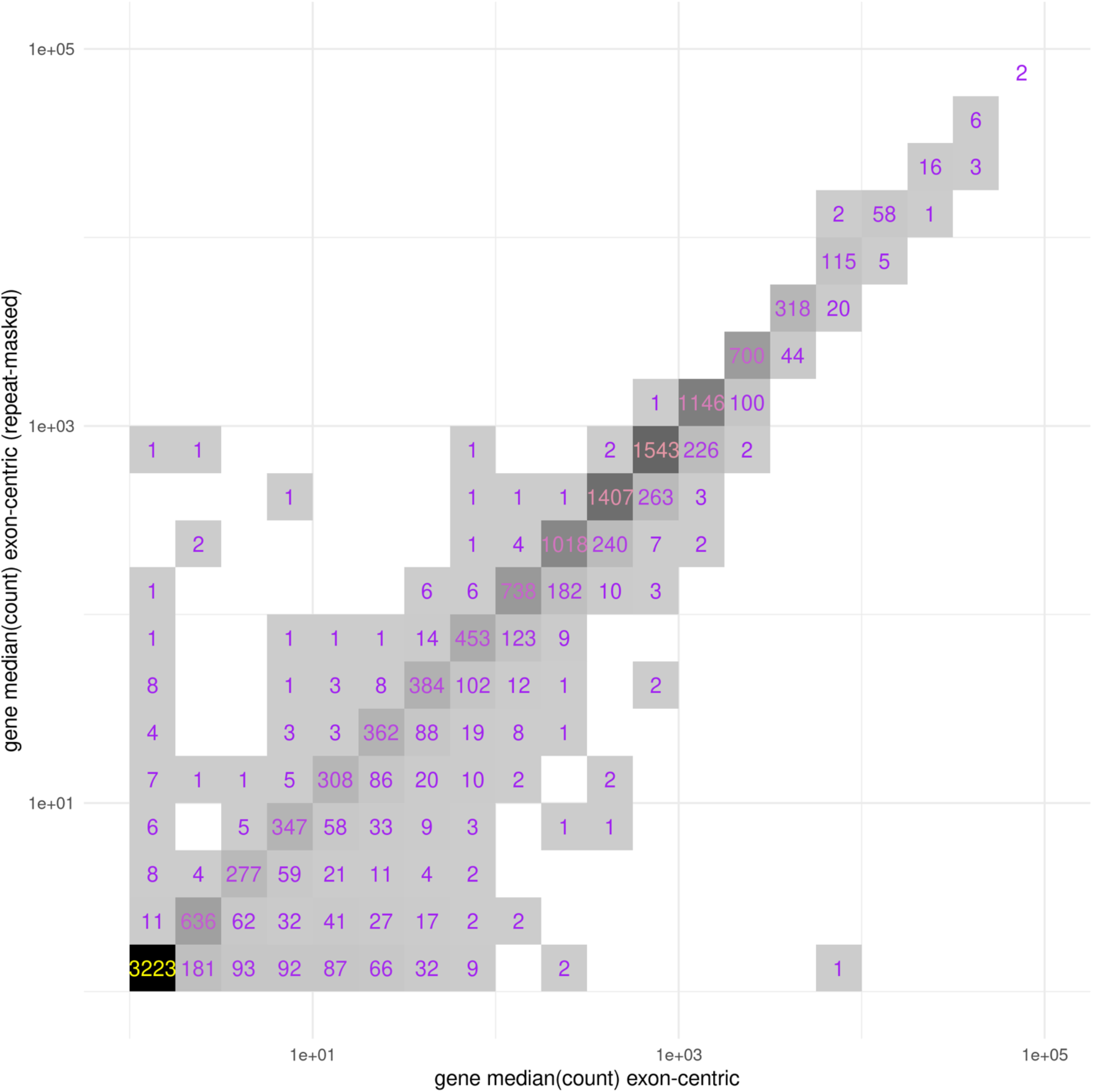
Median estimated read counts per gene for MCF7 models (Table 1) for either exon-centric transcript models (x-axis) or exon-centric transcript models with repeat-masking (y-axis) demonstrates inflation (right shifted bins) of gene counts in the exon-centric data. Median counts were binned in quarter-log increments with a pseudo-count added to every observation to avoid undefined values.

Another way to visualize the effect of repeat masking is to examine its impact on differential gene expression results. In Fig. 7 this effect is demonstrated between exon-centric transcriptome indices with and without repeat masking. Off-axis genes contain reads that are potentially mis-mapped due to repetitive sequence content. Copies of these repetitive sequences may be found throughout the genome assembly, and without repeat masking, reads that map well to these sequences may be incorrectly assigned.

**Fig. 7.**
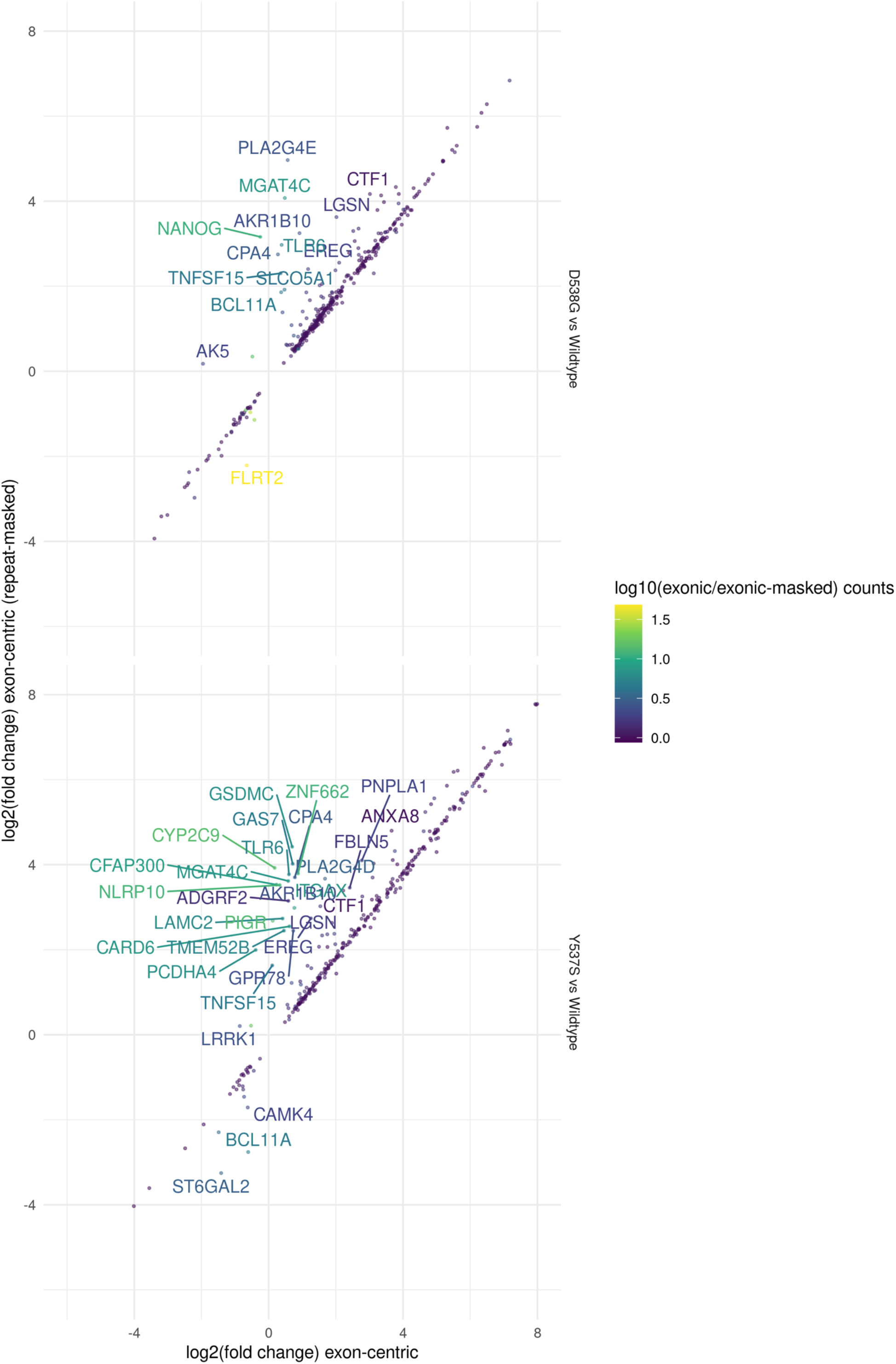
Differentially expressed genes identified in D538G or Y537S vs. Wild-type MCF7 models (Table 1) for either exon-centric transcript models (x-axis) or exon-centric transcript models with repeatmasking (y-axis). Off-axis genes indicate potential mis-mapped reads due to repetitive sequence. DEGs with p < 0.01 are shown. Genes with a greater than 2 fold difference between methods are labeled. Points are colored by the ratio of median gene counts for exon-centric vs exon-centric repeat-masked methods.

Surprisingly, we can achieve almost exactly the same effect as repeat-masking by simply including intronic and intergenic sequences to build a genome-centric transcript index for salmon. This may be due to missing transcripts in our transcriptome model that contain repeats; by including these repeats in our intronic and intergenic sequences they are no longer assigned to the transcript that did have these repeats. LINE and SINE repeats are commonly expressed in several cancer types [14], and transcript models that contain these transposable elements, especially in 5’ and 3’ untranslated regions (UTRs) could be susceptible to such read mis-mapping. Fig. 8 shows that changes to differentially expressed genes are minimal between the repeat-masked exon-centric indexing method and the genome-centric transcriptome indexing method. By compartmentalizing the entire genome assembly we may reduce the number of spurious mappings arising from repetitive genomic sequences.

**Fig. 8.**
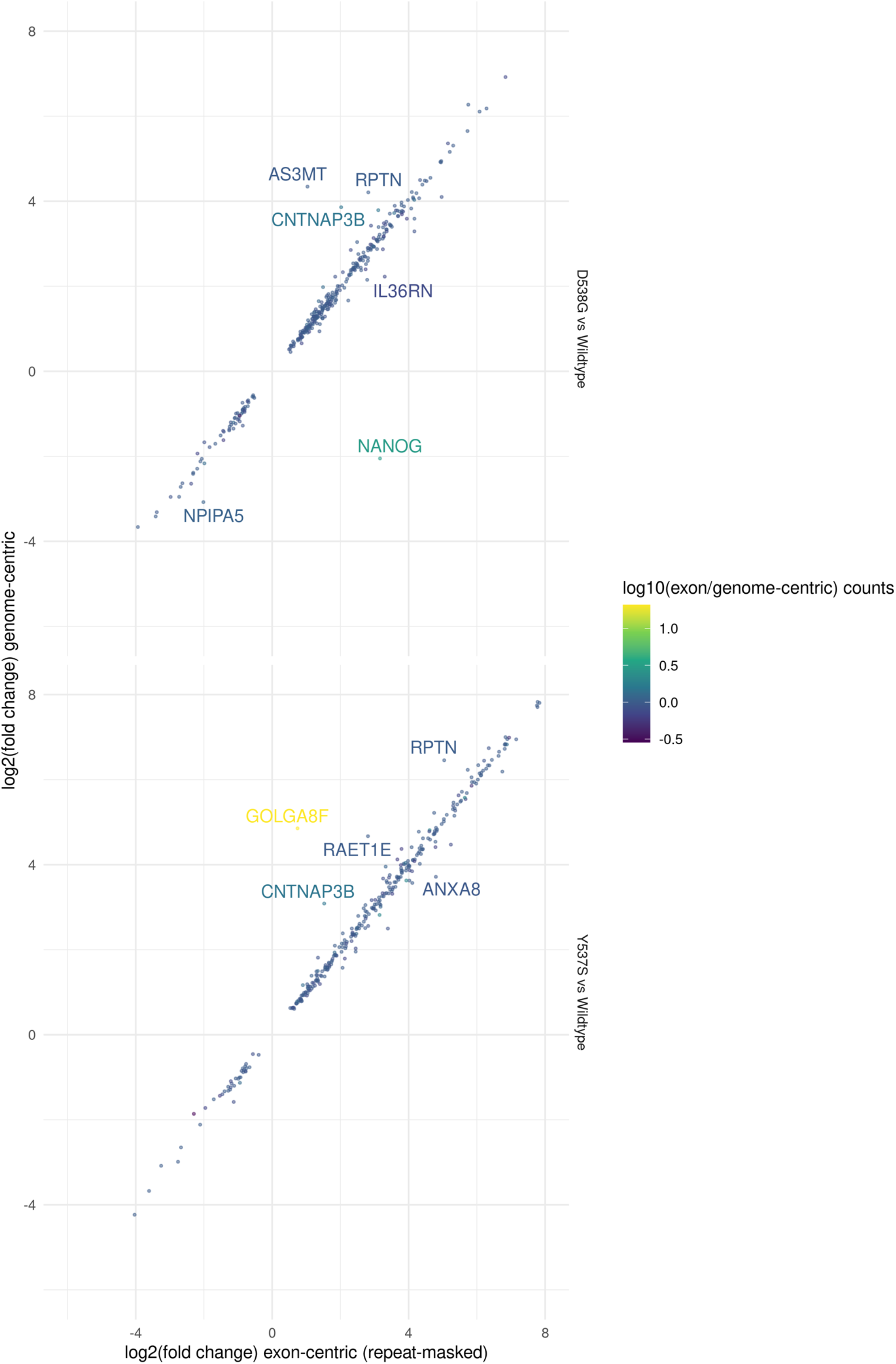
Differentially expressed genes identified in D538G or Y537S vs. Wild-type MCF7 models (Table 1) for either exon-centric transcript models with repeat-masking (x-axis) or genome-centric transcript models (y-axis). Off-axis genes indicate potential mis-mapped reads due to repetitive sequence. DEGs with p < 0.01 are shown. Genes with a greater than 2 fold difference between exon-centric (repeat-masked) and genome-centric methods are labeled. Points are colored by the ratio of median gene counts for exon-centric repeat-masked vs genome-centric methods.

Based on these results we recommend the use of a genome-centric transcriptome index in PISCES.

## Discussion

The motivation for development of PISCES is to enable the efficient, reproducible, and automated analysis of the majority of RNAseq experiments. To this end, the main design goals were ease of installation and use, built-in cluster submission to enable rapid job processing, clarity and reproducibility through use of configuration files, and a design that encourages users to describe experiments using metadata files that can be leveraged at multiple steps in the PISCES workflow. In this way users are encouraged toward reproducible description of their RNAseq analysis.

Traditional alignment-based RNAseq pipelines trade longer run times for more detailed alignment of each sequence read. The gains in computing efficiency acheived by pseudoalignment based transcript quantification come at the cost of this detailed alignment information. This required adaptation of existing RNAseq QC methods that rely on alignment, to keep the total run time of the PISCES pipeline close to the run time of its’ slowest individual component. Integration of external datasets (e.g. TCGA, GTEx, or SRA projects) can be complicated by varying quality control from different sites, data transfer concerns due to disparate (local and remote) data sources, and limitations due to compute times necessary to run traditional tools. PISCES enables large scale integration of datasets, processing of high throughput RNAseq generated from large plate format screens, fast quantification against multiple custom transcriptomes, and fast, reliable reprocessing of libraries whenever new transcriptome annotation or genome assemblies become available.

In addition to gains in computational efficiency, PISCES enables rapid analysis of routine RNAseq experiments, through automated modeling of differential gene expression and generation of clean tables for visualizing gene expression, sample clustering, genetic identity, and QC metrics. We demonstrate a novel genome-centric transcriptome indexing strategy that allows integrated transcript QC measures of intronic and intergenic sequences while also avoiding read mismapping that may occur with exon-centric transcriptome quantification. PISCES is available on GitHub at https://github.com/Novartis/pisces, and on the python package index (PyPI) at https://pypi.org/project/novartis-pisces.

